# Dual-view Guided Context-aware Network for Automated Bone Lesion Segmentation and Quantification in Whole-body SPECT

**DOI:** 10.64898/2026.05.07.723665

**Authors:** Wu Chen, Xiurong Yang, Jing Lu, Miao Miao, Yuan Huang, Shuang Zheng, Chuhan Zhang, Lixuan Xie, Yunfei Zhang

## Abstract

Whole-body SPECT bone scintigraphy reflects skeletal metabolic activity throughout the body and plays an indispensable role in the screening, treatment evaluation, and prognostic assessment of bone metastases in tumors. However, the automatic detection and segmentation of hypermetabolic bone lesions remain challenging due to low contrast, limited spatial resolution, and complex lesion distributions. In this study, we proposed Bone-Segnet, a dual-view guided automatic segmentation network for hypermetabolic bone lesions that integrated multi-scale feature modeling, global context modeling, and view-conditioned modulation. Pixel-level annotated anterior and posterior whole-body bone scintigraphy images were used for model training and prediction. The proposed network enhanced the recognition of low-contrast and small-scale lesions through small-lesion enhancement and multi-scale contextual modeling. A Transformer module was further introduced to strengthen global feature representation, while cross-view collaborative modeling was achieved by incorporating the complementary characteristics of anterior and posterior imaging. Experimental results demonstrated that the proposed method outperformed existing approaches across multiple evaluation metrics, with the Dice score improving from 0.7440 to 0.8750, indicating a substantial improvement in segmentation performance. Further quantitative analysis based on the segmentation results revealed significant differences among disease types in lesion count, pixel burden, and spatial distribution patterns, reflecting the heterogeneity of disease-related skeletal metabolic activity. Overall, the proposed method improved automatic lesion segmentation performance and enabled quantitative analysis of lesion burden and spatial distribution patterns, providing objective data support for the assessment of related diseases. Index Terms—Whole-body SPECT, bone lesion segmentation, dual-view modeling, quantitative analysis.

## Introduction

Whole-body SPECT (Single photon emission computed tomography, SPECT) bone scintigraphy is a routine imaging technique in nuclear medicine for evaluating bone metabolism. After intravenous injection of the radiotracer technetium-99m methylene diphosphonate (99mTc-MDP), its distribution in the whole skeleton reflects overall bone metabolic activity^[1–3]^. Bone metastases typically present as focal increases in bone metabolic activity, and whole-body SPECT bone scintigraphy is important for their detection, prognosis assessment, and treatment evaluation^[4, 5]^. Compared with regional imaging, a single scan covers the entire skeletal system. It provides global information, including lesion number, distribution, and overall burden. This makes it highly valuable in the clinical management of patients with malignant tumors^[5]^.

Whole-body SPECT bone images have low contrast and limited spatial resolution. Clinical interpretation mainly relies on visual inspection of anterior and posterior images. Physicians review the images one by one and perform a comprehensive analysis to identify abnormal uptake and assess distribution patterns^[6]^. Because these images cover a large area, lesion number and distribution are often complex. This is especially true in cases with multiple or suspicious lesions. Manual interpretation is time-consuming and highly dependent on the reader’s experience^[7–9]^. Therefore, developing automatic detection and segmentation methods for whole-body bone scintigraphy is important. It can improve efficiency and enhance the stability of analysis results.

In recent years, many studies have explored automatic detection and segmentation of hypermetabolic bone lesions in bone scintigraphy^[10–18]^. Early methods were mainly based on rule-based strategies, such as thresholding and region growing^[19, 20]^. However, these methods were sensitive to image quality and often achieved limited accuracy. With the development of deep learning, medical image segmentation has advanced rapidly^[21–29]^. Convolutional neural network (CNN) based methods became the mainstream^[30–34]^. These methods used multi-layer convolution to extract and fuse features. They enabled end-to-end learning and showed advantages in representing complex structures. On this basis, further improvements focused on context modeling and structural constraints to enhance performance. For example, Ma et al.^[17]^ introduced an attention mechanism to improve global context modeling. This enhanced the representation of lesion-related features in bone images. However, under complex backgrounds and multiple lesions, the overall segmentation accuracy remained limited. Overall, existing methods still rely mainly on local convolution operations. In whole-body bone images with a large field of view, their ability to model long-range spatial dependencies is limited. As a result, they cannot fully utilize the global distribution of lesions. In addition, small or low-uptake lesions are often overlooked during feature extraction. Uneven image intensity makes lesion boundaries difficult to define. Physiological high-uptake regions may also introduce false positives. These issues indicate that current methods still need further improvement in segmentation performance and robustness under complex conditions.

In this study, to address the limited segmentation performance of hypermetabolic bone lesions in whole-body bone scintigraphy under low-contrast backgrounds and multi-lesion distributions, we developed Bone-Segnet (Bone segmentation network), a segmentation network that integrates multi-scale feature modeling, global context modeling, and a dual-view conditional modulation mechanism. Pixel-level annotated anterior and posterior whole-body bone scintigraphy images were used to construct the training and evaluation datasets. The proposed network incorporated small-lesion enhancement and multi-scale contextual modeling to improve the detection of low-contrast and small targets. A Transformer module was employed to capture global context, while a view-conditioned modulation mechanism leveraged dual-view information for cross-view feature modeling. Experimental results showed that the proposed method outperformed existing approaches across quantitative evaluation metrics, with the Dice score improving from 0.7440 to 0.8750. Quantitative analysis further revealed that the average lesion number ranged from 4.66 to 11.11 per patient, and the average pixel count per lesion ranged from 60.42 to 125.68, indicating disease-specific differences in bone metabolic activity and lesion burden. Overall, the method improved segmentation performance and enabled quantitative characterization of whole-body lesion burden and spatial distribution patterns, providing a reliable basis for the accurate assessment of related diseases.

## Methods

### Dataset acquisition

A retrospective analysis was conducted on 932 patients with hypermetabolic bone lesions identified on whole-body bone scintigraphy between September 1, 2024 and December 31, 2025. This study was approved by the Institutional Review Board of the General Hospital of Central Theater Command (No.[2026]041-01). The requirement for informed consent was waived. All patient data were anonymized prior to analysis.

Whole-body bone imaging was performed 3-4 hours after intravenous injection of 740-1110 MBq technetium-99m methylene diphosphonate (99mTc-MDP, Wuhan Atom High-Tech Pharmaceutical Co., Ltd.). Imaging was acquired using a single photon emission computed tomography (SPECT) system equipped with a low-energy high-resolution parallel-hole collimator (GE Discovery NM 630 dual-head system). A continuous whole-body scanning mode was used, with a scanning speed of approximately 16 cm/min and a matrix size of 256 × 1024. The images were 16-bit DICOM data, including anterior and posterior views. Based on this dataset, the segmentation annotations of hypermetabolic bone lesions were performed by experienced nuclear medicine physicians.

### Training Data Generation

Whole-body bone scintigraphy includes anterior and posterior views. First, the original images were preprocessed, including size normalization and intensity normalization, to ensure consistency for annotation and model training. Then, experienced nuclear medicine physicians performed pixel-level annotation of hypermetabolic bone lesions and generated corresponding segmentation masks. Multiple annotators independently labeled the images, followed by cross-review. Disagreements were discussed and resolved to improve accuracy and consistency.

To meet the input requirements of deep learning models and enhance the learning of local lesion features, the annotated images and masks were further cropped into 256 × 256 patches, while maintaining a one-to-one correspondence between images and labels. Finally, paired anterior and posterior image patches with lesion annotations were constructed as the training dataset and split into training and validation sets. This dataset supported model training and performance evaluation, and also provided a reliable basis for subsequent quantitative analysis of lesion distribution patterns.

### Bone Lesion Segmentation Network

In the task of automatic segmentation of hypermetabolic bone lesions, it is important to select a base model with strong multi-scale feature representation and structural flexibility. In this study, MK-UNet^[35]^ was adopted as the backbone network. Based on the classical U-Net^[36],^ the network employs lightweight multi-kernel convolutions to build multi-resolution feature representations and integrates a multi-kernel attention mechanism to progressively refine multi-scale features, enabling robust target extraction across different receptive fields. Compared with the conventional U-Net, MK-UNet shows better adaptability to low-contrast and morphologically diverse targets in complex backgrounds, providing a reliable basis for hypermetabolic bone lesion segmentation in whole-body SPECT images.

However, in whole-body bone scintigraphy, hypermetabolic bone lesions may exhibit wide spatial distribution, variable lesion counts, and diverse morphologies, posing challenges for accurate segmentation. In practice, MK-UNet still has limitations, including incomplete detection of multiple lesions, missed small or low-contrast regions, and insufficient modeling of long-range spatial dependencies. These limitations may compromise the completeness and stability of segmentation results. Therefore, we build upon MK-UNet to enhance the representation of complex lesion distributions and small-scale targets, thereby improving overall segmentation performance.

In this study, we developed a segmentation network for hypermetabolic bone lesions in whole-body bone scintigraphy, named Bone-Segnet (Bone segmentation network). The network used images as the main input and introduced view-conditioned modeling. This encoded anterior or posterior imaging information and modulated feature representation. The input first passed through the encoder for multi-scale feature extraction. Multi-scale feature enhancement was introduced during encoding. In shallow and intermediate layers, small-lesion enhancement and multi-scale context modules were applied. These modules strengthened local details and improved the representation of small lesions. In high-level features, a Transformer module was used for global context modeling. It captured long-range spatial relationships across the whole body. These features were fused in the encoding stage. The fused features were then fed into the decoder. The decoder performed progressive upsampling. Skip connections and attention gates were used for cross-layer fusion. Spatial resolution was gradually restored, and segmentation boundaries were refined. Finally, the segmentation results were generated.

Compared with the baseline, Bone-Segnet introduced several targeted improvements while retaining the multi-kernel encoder-decoder structure. In shallow layers, a small-lesion enhancement module was added. It used parallel multi-kernel convolutions and channel and spatial attention to improve responses to small and low-contrast targets^[37, 38]^. In intermediate layers, a Midcontext refine module was introduced. It combined multi-scale dilated convolutions and attention to enhance local and mid-level context representation^[39]^. In the deepest layer, a Transformer bottleneck was applied for global context modeling. This improved the ability to capture long-range spatial dependencies^[40]^. In addition, view-conditioned embedding and a view condition gate were used to modulate features from anterior and posterior images. This enabled dynamic fusion of cross-view information. In the decoding stage, grouped attention gates, a detail fusion block, and an output refinement head were added. These modules improved boundary reconstruction and preserved fine details. Overall, these improvements enhance the model’s representation of small-scale targets, complex spatial distributions, and cross-view information, thereby improving hypermetabolic bone lesion segmentation performance.

### Feature Enhancement Modules

To improve feature representation under complex conditions in whole-body bone scintigraphy, several feature enhancement modules were introduced into the base network. These modules focused on local detail enhancement, global context modeling, and cross-view information utilization.

The small lesion enhancement module was embedded in the shallow stage of the encoder. It was used to enhance initial features. The input features were first processed by a multi-branch convolution structure. Different branches used convolutions with different kernel sizes and dilated convolutions to capture multi-scale local responses. The features from all branches were then fused. Channel attention and spatial attention were applied to reweight the features and highlight potential hypermetabolic regions. A residual connection was used to combine the enhanced features with the original features. This design improved the representation of small and low-contrast lesions without destroying the original structural information.

The view-conditioned module was introduced in the high-level encoder. In whole-body bone scintigraphy, anterior and posterior images share structural correspondence but differ in signal distribution, posing challenges for segmentation networks with shared parameters and limiting their ability to fully exploit view-specific information. To address this issue, a view-conditioned module is introduced at the high-level stages of the encoder, where view information is incorporated as an explicit prior into the feature learning process. Specifically, discrete view labels are first mapped into continuous embedding vectors and projected to match the channel dimensions of high-level features. The embedded information is then integrated into the feature space through two complementary pathways: additive fusion to inject global semantic context, and a gating mechanism to adaptively modulate feature responses, enabling feature reconstruction under view-conditioned constraints. This design allows the network to maintain shared backbone parameters while gaining view-awareness, thereby enhancing the consistency and complementarity of cross-view feature representations.

### Loss function

To address the sparse foreground, missed detection, and insufficient overlap in hypermetabolic bone lesion segmentation, multiple loss functions were jointly used to optimize the model.

The Focal Tversky Loss (FTL)^[41]^ is based on the Tversky index. It assigns different weights to false positives and false negatives, which helps handle class imbalance. It also increases the model’s focus on missed lesions during optimization. This makes the model more robust when the foreground is small and lesion size varies. The formulation of this loss is given as follows.

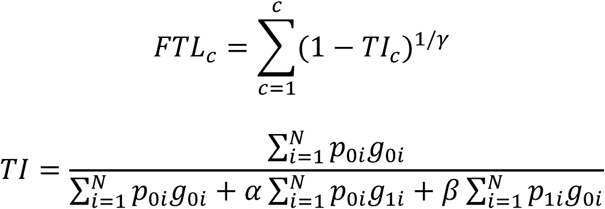

Here, TI denotes the Tversky index, and γ varies in the range [1, 3]. This loss has an adaptive regulation ability for samples with different difficulty levels during optimization. When the Tversky index is high, local errors have a limited effect on the overall loss. When the Tversky index is low, the loss becomes more sensitive to errors, which increases the model’s focus on difficult regions. When γ = 1, the FTL simplifies to the TL.

Where *p*_0*i*_ is the probability of pixel *i* belonging to the foreground class and *p*_1*i*_ is the probability of a pixel belonging to the background class. *g*_0*i*_ is 1 for foreground and 0 for background, whereas *g*_1*i*_ is 1 for background and 0 for foreground. By assigning a higher weight to false negatives, the model’s detection ability was improved. This helped balance precision and recall. Therefore, in this study, β was set greater than α. Specifically, β = 0.70 and α = 0.30 were used to enhance the recognition of hypermetabolic bone regions.

Dice loss is mainly used to address class imbalance in medical image segmentation^[42]^. In such cases, the background occupies a large proportion, while the target region is small. This loss evaluates segmentation performance by measuring the overlap between the prediction and the ground truth. It guides the model to focus more on accurate segmentation of the foreground. To avoid division by zero during computation, a smoothing term is usually added to both the numerator and denominator to improve numerical stability. The formulation is given as follows.

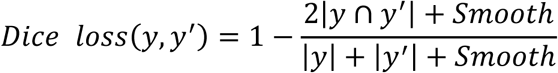

Here, the value of *Smooth* is usually set to 1. The variable *y* denotes the predicted result, and *y*^′^ denotes the ground truth. The term |*y* ∩ *y*^′^| represents the intersection between the prediction and the ground truth, while |*y*| + |*y*^′^| represents the sum of the two sets.

Binary Cross Entropy (BCE) loss is a commonly used loss function for binary classification^[43]^. It measures the difference between predicted probabilities and ground truth labels, and is widely applied in medical image segmentation. This loss applies independent constraints at the pixel level, which helps improve the model’s ability to distinguish target regions. The formulation is given as follows.

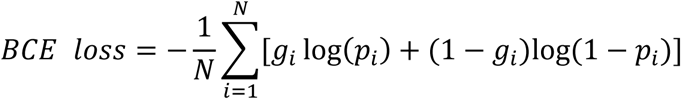

Here, *p*_*i*_ denotes the predicted probability that pixel *i* belongs to the target region, *g*_*i*_ is the corresponding ground truth label, and *N* is the total number of pixels.

This study considered the low foreground proportion, frequent missed detections, and insufficient overlap in hypermetabolic bone lesion segmentation. A composite loss function was used to jointly optimize the network. It consisted of Focal Tversky Loss, Dice Loss, and BCE Loss. The formulation is given as follows.

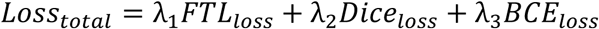

Here, λ_1_, λ_2_, and λ_3_ denote the weight coefficients of each loss term. Based on the characteristics of missed detection and class imbalance in hypermetabolic bone lesion segmentation, empirical weights were assigned to different loss terms. This helped the model achieve a balance among detection ability, region consistency, and pixel-level discrimination.

### Network Training

The network was trained using the PyTorch deep learning framework. The input data consisted of cropped 256 × 256 grayscale images and their corresponding pixel-level annotations, with one-to-one matching based on file names. The input images were normalized before being fed into the model to improve training stability.

During training, data were loaded in a mini-batch manner. The batch size was set according to GPU memory, such as 16 or 32. To improve efficiency and reduce memory usage, automatic mixed precision (AMP) was adopted. Stable training was achieved through dynamic gradient scaling. The model output was a probability map with the same size as the input. After activation, it was used for loss computation. Model optimization was performed using a gradient descent-based method, with an initial learning rate of 3 × 10^⁻5^. A learning rate scheduler was applied during training. The learning rate was adjusted dynamically based on validation performance to promote convergence and avoid local minima.

The dataset was split into training and validation sets. After each epoch, the model was evaluated on the validation set, and the best-performing model was saved. An early stopping strategy was also used. Training was terminated when performance did not improve over several epochs, which helped reduce overfitting.

In addition, random data augmentation was applied during data loading, such as horizontal flipping and random rotation. This improved the model’s robustness to different lesion shapes and complex backgrounds. After training, inference was performed on an independent test set, and the segmentation results were quantitatively evaluated.

The experiments were conducted on a platform with an NVIDIA Quadro RTX 5090 GPU with 32 GB memory, two 32-core CPUs (Intel Xeon Gold 6246R) at 3.39 GHz, and 256 GB RAM. This configuration met the requirements for model training and inference.

### Network Predicting

During the model prediction stage, the size of whole-body bone images is large, so it is difficult to input the entire image into the model directly. Therefore, a sliding-window patch-based prediction strategy was used. This reduced memory usage and improved computational efficiency. Specifically, the image was cropped into patches with a fixed size of 256 × 256. Adjacent patches had overlapping regions to maintain continuity at the boundaries. Each patch was fed into the model for prediction to obtain its segmentation result.

During reconstruction, the predicted patches were stitched back according to their original spatial positions. In overlapping regions, weighted fusion or averaging was applied to combine multiple predictions. This reduced boundary artifacts and discontinuities caused by patching. It also improved the smoothness and consistency of the final segmentation. The reconstructed segmentation map was then used for quantitative analysis and distribution pattern study of hypermetabolic bone lesions.

### Evaluation Metrics

To quantitatively evaluate the segmentation performance of hypermetabolic bone lesions, Precision, Recall, Dice, and Jaccard were used as the main metrics^[7, 8]^. These metrics were calculated based on pixel-level comparison between the predicted results and the corresponding annotations. Precision represents the proportion of correctly predicted lesion pixels among all predicted lesion pixels. Recall represents the proportion of true lesion pixels that were correctly identified, reflecting the detection ability of the model. The Dice coefficient measures the overlap between the predicted region and the ground truth, and reflects the balance between segmentation accuracy and completeness. Jaccard represents the ratio of the intersection to the union between the predicted and ground truth regions, and evaluates the overall consistency of the segmentation. The formulations are given as follows.

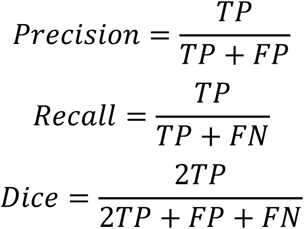

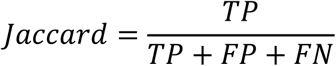

Here, *TP* denotes the number of pixels correctly predicted as hypermetabolic bone lesions. *FP* and *FN* denote the number of falsely predicted lesion pixels and the number of missed true lesion pixels, respectively.

These metrics enable quantitative analysis of the segmentation results. They provide a comprehensive evaluation of the model’s performance in lesion detection. They also offer an objective basis for comparing different methods and for subsequent analysis of distribution patterns.

## Results

### Segmentation Framework of Hypermetabolic Bone Lesions in Whole-Body Bone Scintigraphy

Hypermetabolic bone lesions in whole-body bone scintigraphy show high complexity in spatial distribution and morphology. These lesions are often multiple. They vary greatly in size and shape, and different patterns may appear in the same image. In addition, physiological high uptake and background noise can reduce contrast in some regions. Boundaries may become unclear and can be confused with normal structures or nonspecific uptake. Under these conditions, existing convolutional neural network-based methods still show insufficient detection and incomplete boundary delineation in multi-lesion scenarios. This limits overall segmentation performance.

As shown in Fig. 1(a), typical results of existing networks can be observed. In multi-lesion cases, some small or low-contrast regions are not effectively detected, leading to missed lesions. This problem reduces the overall detection accuracy and also affects the reliability of subsequent quantitative analysis.

**Fig. 1.**
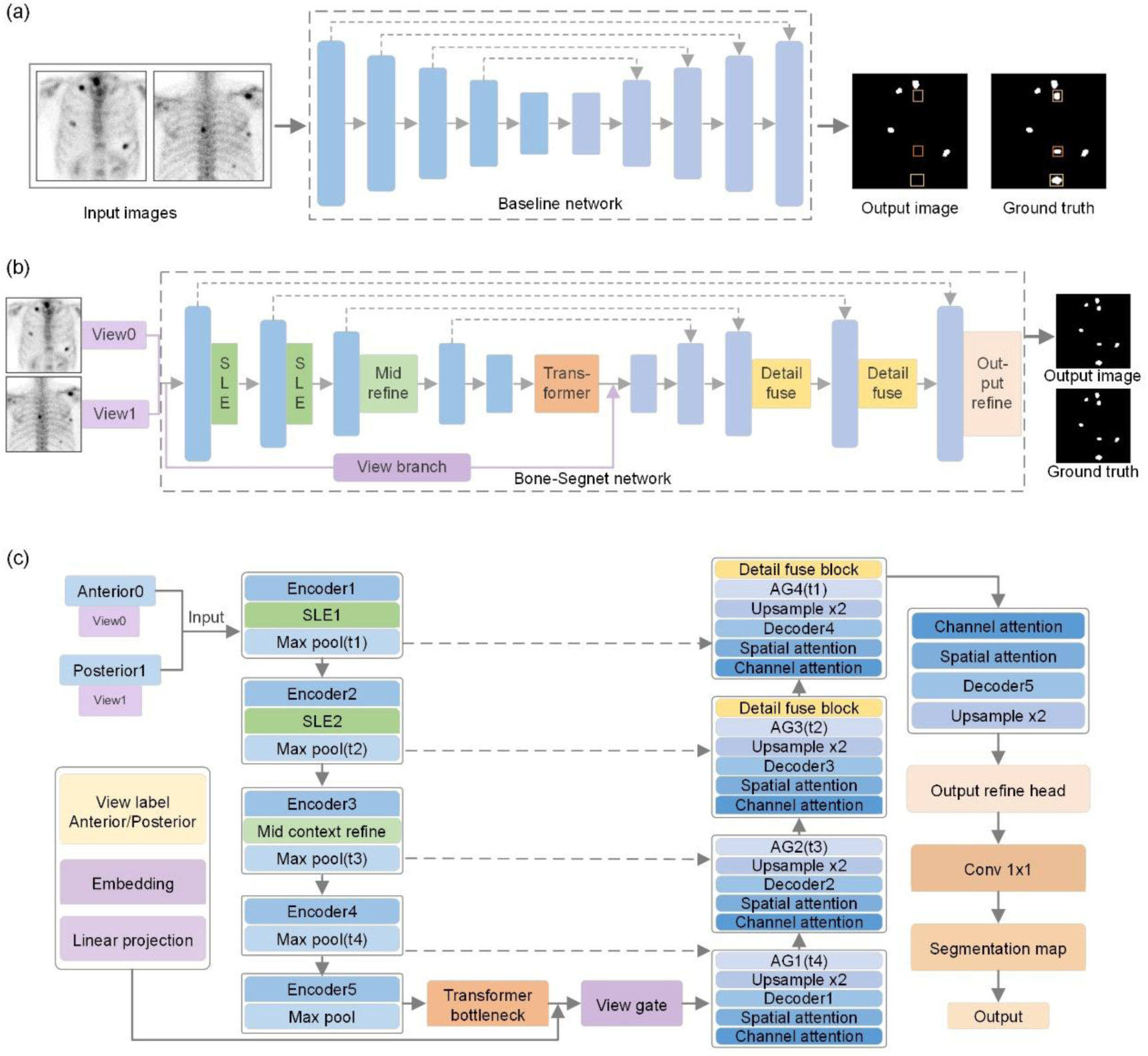
Segmentation framework for hypermetabolic bone lesions in whole-body bone scintigraphy. (a) Limitations of existing neural networks in lesion prediction. (b) Prediction results of the proposed segmentation network. (c) Architecture of the segmentation network for hypermetabolic bone lesions.

To address these issues, a segmentation network named Bone-Segnet was developed based on MK-UNet. It integrates multi-scale feature modeling, global context modeling, and view-conditioned modulation. This design improves segmentation accuracy and stability under complex backgrounds and multi-lesion conditions, as shown in Fig. 1b. The results of Bone-Segnet show more complete detection in multi-lesion regions. Lesions of different sizes are well covered. By introducing multi-scale feature modeling, a Transformer for global context, and view-conditioned modulation, the network improves feature representation for small and low-contrast regions. It also enhances the modeling of long-range spatial relationships across the whole body. As a result, more stable segmentation is achieved under complex conditions.

Fig. 1(c) shows the framework of the proposed segmentation network for hypermetabolic bone lesions. It illustrates the feature extraction and reconstruction process based on an encoder–decoder structure, as well as the collaboration of key modules such as multi-scale feature enhancement, Transformer-based global modeling, and view-conditioned modulation. The encoder extracts multi-scale features layer by layer. In the decoder, skip connections are used for progressive reconstruction and fusion of features.

On this basis, a small lesion enhancement (SLE) module is introduced to strengthen the representation of small lesions. A Transformer module is embedded to enhance global spatial modeling. A ViewGate module is applied to adaptively modulate feature responses using view information. In addition, a Detail Fuse Block is used in the decoding stage to further integrate multi-scale details and improve the representation of fine structures. An Output Refine Head is applied at the final stage to refine the segmentation results. The cooperation of these modules improves the representation of lesions at different scales and enhances segmentation stability under complex backgrounds.

### Data Processing and Segmentation Results of Bone-Segnet

To construct a dataset for model training and evaluation, hypermetabolic bone lesions in whole-body bone scintigraphy were manually annotated. For anterior and posterior images, experienced nuclear medicine physicians performed pixel-level delineation of lesion regions, and the results were saved as binary masks. During annotation, special attention was given to consistency in identifying multiple lesions, small lesions, and low-contrast regions, to ensure accuracy and stability. Some annotation examples are shown in Fig. 2.

**Fig. 2.**
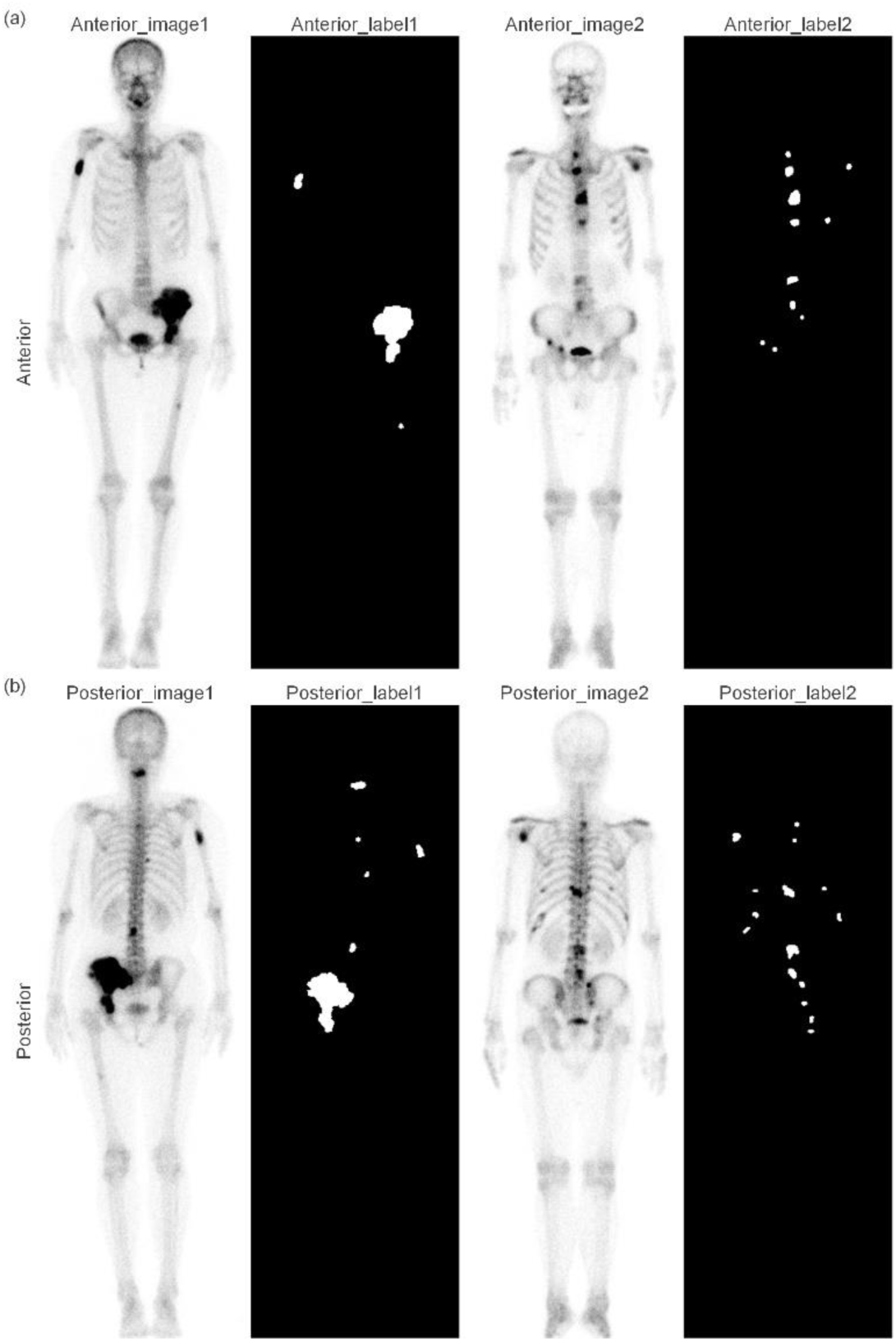
Annotation of hypermetabolic bone lesions in whole-body bone scintigraphy. (a) Anterior images and corresponding annotations. (b) Posterior images and corresponding annotations.

During data preprocessing, the images were cropped to improve local feature learning and to unify the input size. The original whole-body images were divided into patches of 256 × 256 using a sliding window. Each patch had a corresponding annotation. These patches were used to construct the training dataset. During cropping, different anatomical regions and lesions of different sizes were covered as much as possible. This ensured diversity of the training samples. This strategy preserved local structural information and increased the number of samples. It also enhanced the model’s ability to learn multi-scale features, as shown in Fig. 3a.

**Fig. 3.**
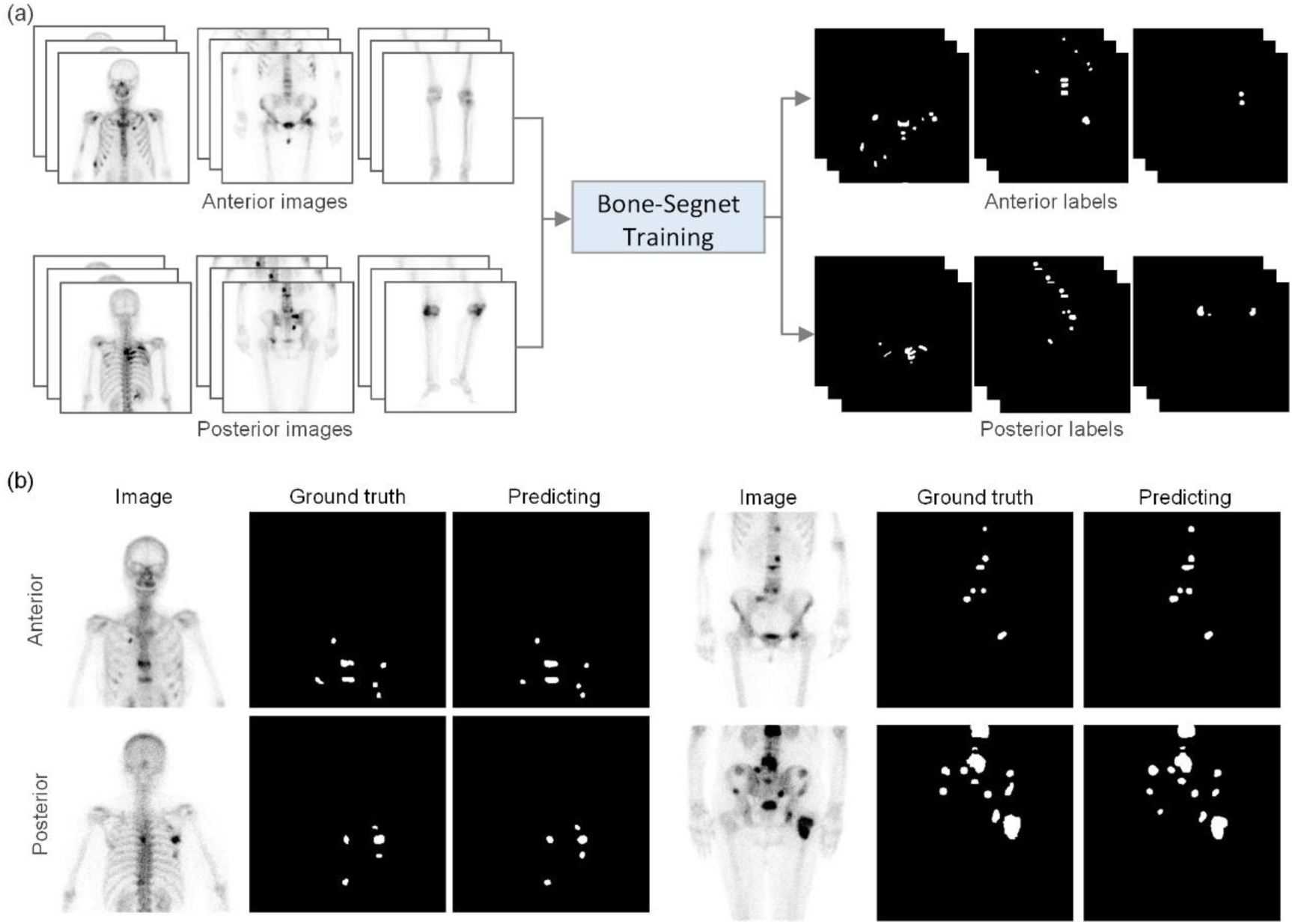
Training and prediction of the Bone-Segnet network. (a) Image cropping and network training. (b) Predicted hypermetabolic lesions in anterior and posterior images by Bone-Segnet.

After training, the Bone-Segnet model was applied to the test data for prediction. The segmentation results of hypermetabolic bone lesions were obtained. As shown in Fig. 3b, the model identified lesions across the whole body and provided relatively complete segmentation for targets of different sizes. In multi-lesion cases, most lesions were successfully detected. The model also maintained reasonable performance in low-contrast regions. Overall, the predictions were consistent with manual annotations, showing good segmentation ability and stability under complex conditions.

### Ablation Study

To systematically evaluate the contribution of each module, an ablation study was designed. Six models, Net1 to Net6, were constructed by progressively adding different components to the baseline network. All models were trained and evaluated on the same dataset with identical settings to ensure fair comparison. Overall, as the Transformer module, view branch, small-lesion enhancement, and detail fusion were gradually introduced, the performance improved on quantitative metrics. This indicates that each module made a positive contribution to lesion segmentation, as shown in Table 1.

**Table 1.** Ablation study results of different network configurations.

| Model ID | #1 | #2 | #3 | #4 | #5 | Precision | Recall | Dice | Jaccard |
| --- | --- | --- | --- | --- | --- | --- | --- | --- | --- |
| Net1 | √ | × | × | × | × | 0.7553 | 0.7877 | 0.7533 | 0.6116 |
| Net2 | √ | √ | × | × | × | 0.7822 | 0.7758 | 0.7637 | 0.6245 |
| Net3 | √ | × | √ | × | × | 0.7952 | 0.7794 | 0.7716 | 0.6345 |
| Net4 | √ | √ | √ | × | × | 0.7849 | 0.8033 | 0.7792 | 0.6450 |
| Net5 | √ | √ | √ | √ | × | 0.8144 | 0.8591 | 0.8277 | 0.7091 |
| Net6 | √ | √ | √ | √ | √ | 0.8502 | 0.9282 | 0.8843 | 0.7937 |

#1: Back-bone network; #2: Transformer module; #3: View branch; #4: Small lesion/ Context enhancement; #5: Detail fusion and output refinement

Starting from the baseline Net1, adding the Transformer module in Net2 increased the Dice score from 0.7533 to 0.7637. This shows that global context helps represent complex structures. When the view branch was further added in Net3, the Dice score increased to 0.7720. This result is higher than Net2 and suggests that joint modeling of anterior and posterior views is effective. When both the Transformer and the view branch were combined in Net4, the Dice reached 0.7792, and Recall increased to 0.8033. This indicates that global modeling and multi-view information are complementary and improve overall lesion detection.

On this basis, adding the small lesion and context enhancement module in Net5 increased Recall from 0.8033 to 0.8591, and Dice further improved to 0.8277. This indicates better detection of small and low-contrast targets. After introducing the detail fusion and output refinement modules in Net6, the performance improved further. Dice reached 0.8843, which is about 13.10% higher than the baseline. Jaccard increased to 0.7937, and Recall rose to 0.9282. These results show that the detail fusion module plays an important role in improving boundary representation and overall segmentation quality. Overall, the modules worked together at different levels. The view branch and small lesion enhancement mainly improved detection, while the Transformer and detail fusion modules enhanced feature representation and boundary accuracy, leading to consistent performance gains.

### Comparison of Results Across Different Segmentation Networks

To validate the effectiveness and applicability of the proposed Bone-Segnet for hypermetabolic bone lesion segmentation in whole-body SPECT, several commonly used models were selected for comparison. These included U-Net^[36]^, U-Net++^[44]^, Attention U-Net^[45]^, ResUNet++^[46]^, and MK-UNet^[35]^. All models were trained and tested under the same experimental settings. The results are shown in Fig. 4. The figure presents segmentation examples and quantitative comparisons across models. A combined qualitative and quantitative evaluation was performed.

**Fig. 4.**
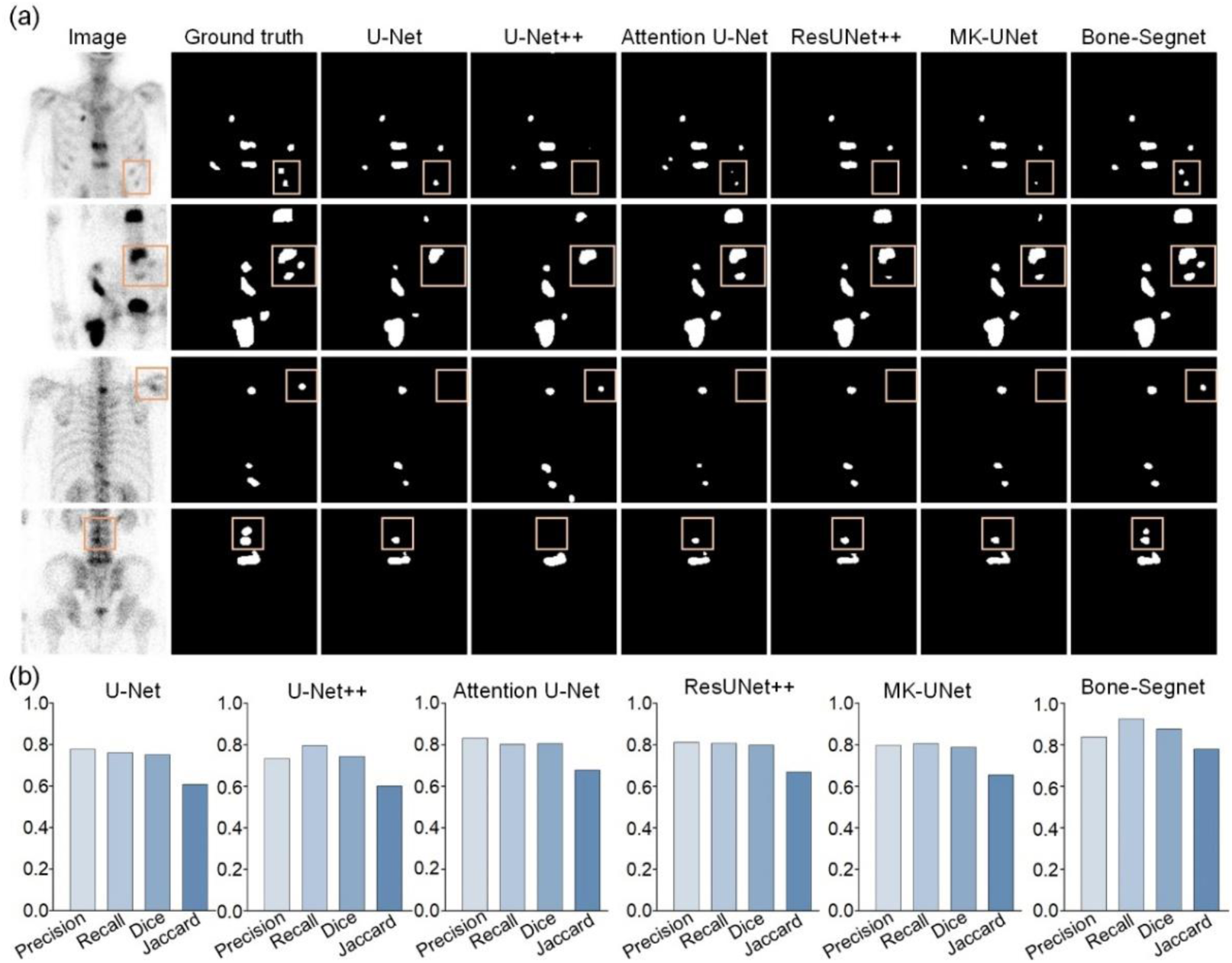
Comparison of different segmentation networks for hypermetabolic bone lesion segmentation in whole-body bone scintigraphy. (a) Segmentation results of different networks. (b) Quantitative evaluation metrics of different networks.

As shown in Fig. 4(a), all comparison models showed similar limitations. Missed detections and incomplete segmentation were observed to varying degrees. These issues were more evident in low-contrast images. The models had limited ability to detect small or weak lesions. In contrast, Bone-Segnet showed better stability and consistency. It achieved more complete coverage of target regions while better preserving lesion integrity. It also performed better in detecting small lesions.

Further analysis was conducted using quantitative metrics, as shown in Fig. 4(b). Compared with other models, Bone-Segnet showed improvements in Precision, Recall, Dice, and Jaccard. Specifically, it achieved the highest Dice score of 0.8750. This represents increases of 12.45%, 13.10%, 6.95%, 7.69%, and 8.69% over U-Net, U-Net++, Attention U-Net, ResUNet++, and MK-UNet, respectively. Overall segmentation performance was improved. These results indicate that global feature modeling and cross-view information improved the model’s adaptability. The model performed better under low-contrast conditions and in multi-lesion scenarios. It also improved the segmentation of small and weak-signal regions. This provides a more stable basis for subsequent quantitative analysis of distribution patterns.

### Quantitative Analysis of Hypermetabolic Bone Lesions

In this study, the proposed Bone-Segnet model was used to automatically segment hypermetabolic bone lesions in whole-body SPECT images. The results were then manually reviewed. Based on this, quantitative analysis of lesion number and pixel burden was performed, as shown in Table 2. The total pixel count of lesions was calculated from the segmentation results and can reflect the relative lesion burden to some extent.

**Table 2.** Distribution of hypermetabolic bone lesions across different diseases.

| Disease type | Patients | Lesion count | Total lesion pixel | Avg. lesions/patient | Avg. pixels /patient | Avg. pixels/lesion |
| --- | --- | --- | --- | --- | --- | --- |
| Lung cancer | 185 | 1,251 | 90,950 | 6.7622 | 491.6216 | 72.7018 |
| Breast cancer | 128 | 596 | 38,267 | 4.6563 | 298.9609 | 64.2064 |
| Prostate cancer | 123 | 1,366 | 171,681 | 11.1057 | 1395.7805 | 125.6816 |
| Nasopharyngeal cancer | 34 | 216 | 15,312 | 6.3529 | 450.3529 | 70.8889 |
| Colon cancer | 33 | 209 | 12,799 | 6.3333 | 387.8485 | 61.2392 |
| Gastric cancer | 24 | 212 | 12,809 | 8.8333 | 533.7083 | 60.4198 |

Among the 932 patients, the six most common disease types were selected for analysis. These included lung malignant tumor, breast malignant tumor, prostate malignant tumor, nasopharyngeal malignant tumor, colon malignant tumor, and gastric malignant tumor. Clear differences were observed in lesion number and burden among diseases. Overall, the average number of lesions ranged from 4.65 to 11.11 per patient. The average pixel count per lesion ranged from 60.42 to 125.68. These results indicate notable differences in bone metabolic activity and lesion scale across diseases. Among them, lung malignant tumors showed more than one thousand lesions in total and a high pixel burden. The average lesion number was 6.7622 per patient, and the total pixel burden reached 90,950, indicating a relatively wide extent of bone involvement. In contrast, breast malignant tumors showed an average of 4.6563 lesions per patient, with an average of 64.2064 pixels per lesion, suggesting fewer lesions and smaller lesion size overall.

Based on the analysis of lesion pixel burden, the number of lesions in different anatomical regions was further analyzed across diseases, as shown in Fig. 5. Structured information extracted from segmentation results enabled more efficient and consistent quantification of multi-site lesions. The results showed a clear imbalance in spatial distribution. The spine and thorax were the most frequently involved regions. Most diseases had higher lesion counts in these areas than in others. For example, lung cancer showed the highest counts in the spine 197 lesions and thorax 168 lesions. Breast cancer and prostate cancer showed similar patterns, with spinal lesion counts reaching 110 lesions and 217 lesions, respectively. These findings suggest that the axial skeleton is a common site of abnormal bone metabolism.

**Fig. 5.**
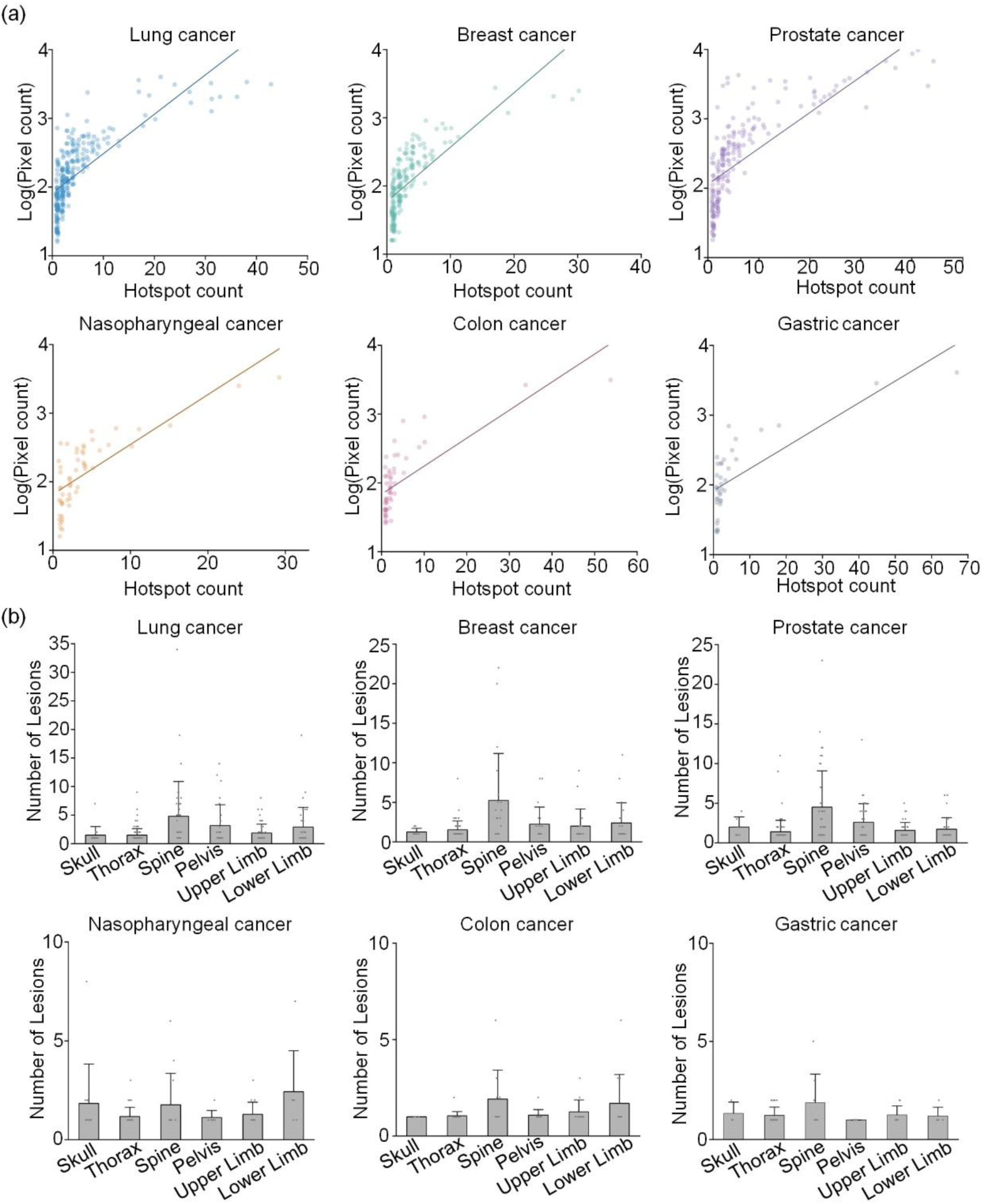
Lesion number and spatial distribution of hypermetabolic bone lesions across different diseases. (a) Scatter plot and linear fitting of total lesion pixel count versus total lesion number for six diseases. (b) Distribution of lesion counts in different anatomical regions for six diseases.

Further analysis showed differences in the pelvis and limb regions among diseases. Prostate-related diseases showed more involvement in the pelvis. Lung cancer also showed relatively high lesion counts in both upper and lower limbs, indicating a wider distribution. In contrast, nasopharyngeal cancer and colon cancer had fewer lesions overall, with a more even distribution across regions. These results indicate clear differences in spatial distribution patterns among diseases. The axial skeleton is the main site of involvement, while peripheral distribution shows disease-specific characteristics.

## Discussion

Whole-body bone scintigraphy has limited spatial resolution and low contrast. Under multi-lesion and complex background conditions, detection and segmentation of hypermetabolic bone lesions remain challenging. Clinical interpretation relies heavily on experience and lacks objective quantification. To address these issues, this study developed a segmentation network named Bone-Segnet. It integrates multi-scale feature modeling, global context modeling, and view-conditioned modulation. The method achieved automatic segmentation of hypermetabolic bone lesions. Experimental results showed that it outperformed existing methods on multiple metrics. It demonstrated good accuracy and stability, and enabled effective segmentation under complex conditions.

The strong performance mainly resulted from the combination of multi-level feature modeling strategies. First, small-lesion enhancement and multi-scale context modeling were introduced in shallow and intermediate layers. This improved the representation of small and low-contrast lesions. Second, a Transformer module was applied in high-level features to model global context. This improved the ability to capture long-range spatial dependencies and enhanced detection in multi-lesion scenarios. In addition, a view-conditioned modulation mechanism was introduced based on anterior and posterior imaging. This enabled cross-view feature interaction and improved consistency and stability. These designs balanced local detail modeling and global feature representation, and improved segmentation under complex conditions.

Based on reliable segmentation results, quantitative characterization of lesions was further performed. Compared with traditional qualitative interpretation, this approach provided objective statistics of lesion number and pixel burden. It better reflected differences in metabolic activity and lesion scale. The results showed variations among diseases, suggesting certain disease-specific patterns. This method improved objectivity and consistency, and provided quantitative support for disease evaluation and treatment monitoring.

Overall, this study not only achieved automatic segmentation of hypermetabolic bone lesions but also enabled a transition from qualitative to quantitative analysis in whole-body bone scintigraphy. The quantitative indicators derived from segmentation provided a basis for analyzing lesion burden and spatial distribution. This helps improve the quantitative level of bone imaging analysis and supports clinical applications.

## Conclusion

This study developed a segmentation network named Bone-Segnet. It integrates multi-scale feature modeling, global context modeling, and view-conditioned modulation. The method achieved automatic segmentation and quantitative analysis of hypermetabolic bone lesions in whole-body bone scintigraphy. Experimental results showed that it outperformed existing methods on multiple metrics. The Dice score improved from 0.7440 to 0.8750, indicating enhanced segmentation performance. Quantitative analysis based on the segmentation results showed differences in lesion number and pixel burden among diseases. These findings indicate that the method not only improves segmentation accuracy but also provides a reliable basis for quantitative evaluation of hypermetabolic bone lesions.

## Appendix

The source code is available at https://github.com/chenwu-mc/Bone-Segnet.

## Acknowledgments

We would like to thank the Department of Nuclear Medicine, General Hospital of Central Theatre Command, Wuhan, China. We also acknowledge the School of Medicine, Jianghan University, Wuhan, China, Hubei Key Laboratory of Cognitive and Affective Disorders, and the Hubei Provincial Demonstration Center for Experimental Medicine Education, School of Medicine, Jianghan University, Wuhan, China.

## Data availability

Data underlying the results presented in this paper are not publicly available at this time, but may be obtained from the authors upon reasonable request.

## References

[1] Van den Wyngaert T, Strobel K, Kampen W U, et al. The EANM practice guidelines for bone scintigraphy[J]. European journal of nuclear medicine and molecular imaging, 2016, 43(9): 1723–1738.

[2] Bombardieri E, Aktolun C, Baum R P, et al. Bone scintigraphy: procedure guidelines for tumour imaging[J]. European journal of nuclear medicine and molecular imaging, 2003, 30(12): B99–B106.

[3] Buckley O, O’Keeffe S, Geoghegan T, et al. 99mTc bone scintigraphy superscans: a review[J]. Nuclear medicine communications, 2007, 28(7): 521–527.

[4] Macedo F, Ladeira K, Pinho F, et al. Bone metastases: an overview. Oncology Reviews, 2017, 11(1): 321

[5] Tsukamoto S, Kido A, Tanaka Y, et al. Current overview of treatment for metastatic bone disease. Current Oncology, 2021, 28(5): 3347–3372

[6] Magdy O, Elaziz M A, Dahou A, et al. Bone scintigraphy based on deep learning model and modified growth optimizer[J]. Scientific Reports, 2024, 14(1): 25627.

[7] Huang K, Huang S, Chen G, et al. An end-to-end multi-task system of automatic lesion detection and anatomical localization in whole-body bone scintigraphy by deep learning. Bioinformatics, 2023, 39(1): 753

[8] Huang Z, Pu X, Tang G, et al. BS-80K: The first large open-access dataset of bone scan images. Computers in Biology and Medicine, 2022, 151: 106221

[9] Wang Y, Lin Q, Zhao S, et al. Automated diagnosis of bone metastasis by classifying bone scintigrams using a self-defined deep learning model. Current Medical Imaging, 2024, 20(1): e15734056281578

[10] Apiparakoon T, Rakratchatakul N, Chantadisai M, et al. MaligNet: semisupervised learning for bone lesion instance segmentation using bone scintigraphy. IEEE Access, 2020, 8: 27047–27066

[11] Lin Q, Gao R, Luo M, et al. Semi-supervised segmentation of metastasis lesions in bone scan images. Frontiers in Molecular Biosciences, 2022, 9: 956720

[12] He Y, Lin Q, Man Z, et al. A Two-Stage CNN-Based Method for Enhanced Metastasis Segmentation in SPECT Bone Scans. International Journal of Intelligent Systems, 2025, 2025(1): 3135835

[13] Wu T, Luo R, Lin H, et al. Research on focal segmentation of bone scan based on Swin Transformer. in: International Conference on Computer Vision, Image and Deep Learning (CVIDL). Proceedings of the IEEE, 2023: 426–430

[14] Sitaba A Y, Rachmawati E, Sulistiyo M D. Investigating convolution-attention model for bone scan image segmentation. in: International Conference on Information Technology, Computer, and Electrical Engineering (ICITACEE). Proceedings of the IEEE, 2023: 344–349

[15] Magdy O, Elaziz M A, Dahou A, et al. Bone scintigraphy based on deep learning model and modified growth optimizer. Scientific Reports, 2024, 14(1): 25627

[16] Xie A, Lin Q, He Y, et al. Metastasis lesion segmentation from bone scintigrams using encoder-decoder architecture model with multi-attention and multi-scale learning. Quantitative Imaging in Medicine and Surgery, 2025, 15(1): 689–708

[17] Ma X, Lin Q, Guo S, et al. Multimodal data-driven segmentation of bone metastasis lesions in SPECT bone scans using deep learning. Current Medical Imaging, 2024, 20(1): e15734056324977

[18] Chen Y Y, Yu P N, Lai Y C, et al. Bone metastases lesion segmentation on breast cancer bone scan images with negative sample training. Diagnostics, 2023, 13(19): 3042

[19] Bhargavi K, Jyothi S. A survey on threshold based segmentation technique in image processing[J]. International Journal of Innovative Research and Development, 2014, 3(12): 234–239.

[20] Preetha M M S J, Suresh L P, Bosco M J. Image segmentation using seeded region growing[C]. 2012 International Conference on Computing, Electronics and Electrical Technologies (ICCEET). IEEE, 2012: 576–583.

[21] Biswas M, Kuppili V, Saba L, et al. State-of-the-art review on deep learning in medical imaging[J]. Frontiers in Bioscience-Landmark, 2019, 24(3): 380–406.

[22] Garg S, Singh P. State-of-the-art review of deep learning for medical image analysis[C]. 2020 3rd International Conference on Intelligent Sustainable Systems (ICISS). IEEE, 2020: 421–427.

[23] Iglesias J E, Sabuncu M R. Multi-atlas segmentation of biomedical images: a survey. Medical Image Analysis, 2015, 24(1): 205–219

[24] Huang Z, Wang L, Xu L. DRA-Net: Medical image segmentation based on adaptive feature extraction and region-level information fusion[J]. Scientific Reports, 2024, 14(1): 9714.

[25] Kherde T C, Baraskar T. A comprehensive survey of image segmentation for medical images[C]. 2024 4th International Conference on Sustainable Expert Systems (ICSES). IEEE, 2024: 1137–1144.

[26] Mienye I D, Swart T G, Obaido G, et al. Deep convolutional neural networks in medical image analysis: A review[J]. Information, 2025, 16(3): 195.

[27] Yu H, Yang L T, Zhang Q, et al. Convolutional neural networks for medical image analysis: state-of-the-art, comparisons, improvement and perspectives[J]. Neurocomputing, 2021, 444: 92–110.

[28] Sinha A, Dolz J. Multi-scale self-guided attention for medical image segmentation[J]. IEEE journal of biomedical and health informatics, 2020, 25(1): 121–130.

[29] Wu M, Liu T, Dai X, et al. Hmda: A hybrid model with multi-scale deformable attention for medical image segmentation[J]. IEEE Journal of Biomedical and Health Informatics, 2024, 29(2): 1243–1255.

[30] Yao W, Bai J, Liao W, et al. From cnn to transformer: A review of medical image segmentation models[J]. Journal of Imaging Informatics in Medicine, 2024, 37(4): 1529–1547.

[31] Zhang Z, Wu H, Zhao H, et al. A novel deep learning model for medical image segmentation with convolutional neural network and transformer[J]. Interdisciplinary Sciences: Computational Life Sciences, 2023, 15(4): 663–677.

[32] Mohapatra S, Swarnkar T, Das J. Deep convolutional neural network in medical image processing[M]. Handbook of deep learning in biomedical engineering. Academic Press, 2021: 25–60.

[33] Zhang Y, Wu P, Chen S, et al. FCE-Net: a fast image contrast enhancement method based on deep learning for biomedical optical images[J]. Biomedical optics express, 2022, 13(6): 3521–3534.

[34] Chen W, Liao M, Bao S, et al. A hierarchically annotated dataset drives tangled filament recognition in digital neuron reconstruction[J]. Patterns, 2024, 5(8).

[35] Rahman M M, Marculescu R. Mk-unet: Multi-kernel lightweight cnn for medical image segmentation[C]. Proceedings of the IEEE/CVF International Conference on Computer Vision. 2025: 1042–1051.

[36] Ronneberger O, Fischer P, Brox T. U-net: Convolutional networks for biomedical image segmentation[C]. International Conference on Medical image computing and computer-assisted intervention. Cham: Springer international publishing, 2015: 234–241.

[37] Kou R, Wang C, Peng Z, et al. Infrared small target segmentation networks: A survey[J]. Pattern recognition, 2023, 143: 109788.

[38] Woo S, Park J, Lee J Y, et al. Cbam: Convolutional block attention module[C]//Proceedings of the European conference on computer vision (ECCV). 2018: 3–19.

[39] Chen L C, Zhu Y, Papandreou G, et al. Encoder-decoder with atrous separable convolution for semantic image segmentation[C]. Proceedings of the European conference on computer vision (ECCV). 2018: 801–818.

[40] A. Vaswani, N. Shazeer, N. Parmar, et al. Attention is all you need. in: Advances in Neural Information Processing Systems (NeurIPS), Long Beach, California, USA, 4-9 Dec. 2017, MIT Press, 2017: 6000–6010

[41] Abraham N, Khan N M. A novel focal tversky loss function with improved attention u-net for lesion segmentation[C]//2019 IEEE 16th international symposium on biomedical imaging (ISBI 2019). IEEE, 2019: 683–687.

[42] Milletari F, Navab N, Ahmadi S A. V-net: Fully convolutional neural networks for volumetric medical image segmentation[C]. 2016 fourth international conference on 3D vision (3DV). Ieee, 2016: 565–571.

[43] Li Q, Jia X, Zhou J, et al. Rediscovering bce loss for uniform classification[J]. arXiv preprint arXiv:2403.07289, 2024.

[44] Zhou Z, Rahman Siddiquee M M, Tajbakhsh N, et al. Unet++: A nested u-net architecture for medical image segmentation[C]. International workshop on deep learning in medical image analysis. Cham: Springer International Publishing, 2018: 3–11.

[45] Oktay O, Schlemper J, Folgoc L L, et al. Attention u-net: learning where to look for the pancreas. arXiv preprint arXiv:1804.03999, 2018

[46] Jha D, Smedsrud P H, Riegler M A, et al. Resunet++: An advanced architecture for medical image segmentation[C]. 2019 IEEE international symposium on multimedia (ISM). IEEE, 2019: 225–2255.

